# Physiological Culture Modulates the Cargo and Wound-Healing Potential of MSC Extracellular Vesicles

**DOI:** 10.64898/2026.11.25.769132

**Authors:** Julia Monola, Alisa Ottila, Nikolas Heikkilä, Chris S. Pridgeon, Riina Harjumäki

## Abstract

This study demonstrates that animal-free 3D culture and long-term hypoxic conditioning modulate mesenchymal stromal cell-derived extracellular vesicle (MSC-EV) production and cargo composition without compromising MSC identity. Mimicking the in vivo microenvironment through hypoxic 3D culture increased MSC-EV yield and altered EV molecular composition, highlighting the potential of culture engineering as a strategy to optimise MSC-EV manufacturing. While 3D culture exerted the greatest effect on both cellular and MSC-EV proteomes, increasing protein abundance and complexity associated with translational and peptide biosynthesis pathways, hypoxic conditioning further enhanced the functional activity of 3D-derived MSC-EVs in an in vitro wound-healing model. These findings suggest that even subtle alterations in MSC-EV cargo can influence biological activity. Collectively, our results demonstrate that culture configuration and oxygen tension are important process variables that affect MSC-EV yield, composition, and functionality. This work highlights the importance of integrating physiologically relevant culture conditions into MSC-EV production strategies and provides insights for the development of scalable and clinically translatable MSC-EV therapeutics for wound healing and regenerative medicine.

**Significance statement:** Mesenchymal stromal cell-derived extracellular vesicles are promising cell-free therapeutics for regenerative medicine, but their clinical translation requires scalable manufacturing strategies. This study demonstrates that culture conditions can substantially influence extracellular vesicle production, molecular composition, and biological activity. The findings identify oxygen tension and three-dimensional culture configuration as important determinants of product quality and therapeutic performance, providing insights for the development of clinically translatable regenerative therapies.

**Graphical abstract:** 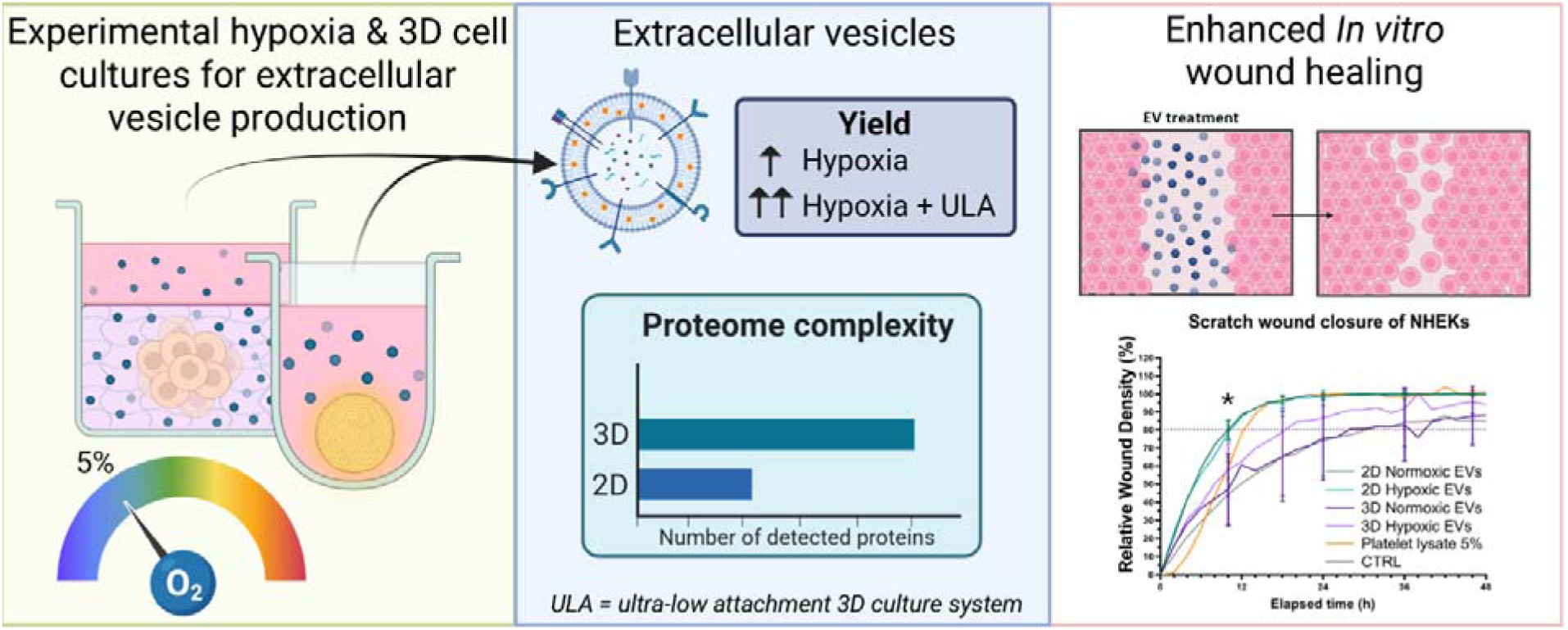

## INTRODUCTION

Mesenchymal stromal cells (MSCs) have been extensively studied for regenerative medicine applications due to their ability to support tissue repair in multiple organs, including neural, cardiac, hepatic, and skin tissues ^1–5^. The therapeutic effects of MSCs are mainly mediated by paracrine signalling through the secretion of growth factors, cytokines, chemokines, and extracellular vesicles (EVs) ^6^. Among these, EVs transfer bioactive cargo, including proteins, lipids, and nucleic acids, to recipient cells, thereby influencing diverse cellular processes. EVs from MSCs retain many of the therapeutic properties of their parental cells while avoiding several safety concerns associated with live-cell therapies, including immunogenicity, tumorigenicity, poor post-transplantation cell survival and logistical challenges making the appealing therapeutic agents ^7–9^.

As cellular phenotype is influenced by the microenvironment, culture conditions alter MSC characteristics and the composition and biological activity of secreted EVs, affecting their molecular cargo and therapeutic efficacy ^10–12^. Therefore, identifying the optimal culture conditions for production of therapeutically effective MSC-EVs is crucial ^10^. In addition to EV composition, the production of EVs in sufficient quantities for clinical applications must be considered when designing culture systems. Consequently, optimising culture conditions to improve both EV yield and biological activity in combination with animal component-free culture is essential for the successful translation of MSC EV-based therapies.

To improve the therapeutic potential and clinical translation of MSC-EVs for regenerative medicine, advanced culture systems that closely mimic the *in vivo* microenvironment have been investigated, including hypoxic conditioning and 3D culture models. By influencing MSC phenotype and function, these techniques can alter EV cargo composition and enhance EV yield, improving their therapeutic, translational, and regenerative potency ^13–19^. In MSCs, 3D culture, which helps to reproduce cell-cell interactions, enhances the expression of pluripotency associated genes ^20–24^, promotes angiogenic properties ^25,26^, and improves differentiation potential ^27^. In addition, 3D culture can increase EV yield and enhance the regenerative efficacy of MSC-EVs^13,22,28–30^.

Oxygen tension *in vivo* varies across tissues, with MSCs residing in naturally hypoxic niches such as the bone marrow, adipose tissue, and umbilical cord, where oxygen levels are typically in the range of 5–7% compared with ∼21% O in ambient air ^31^. To more closely replicate these conditions, hypoxic culture has been used to preserve MSC phenotype and functionality, ultimately improving the therapeutic potential of MSCs and their EVs ^32^. Hypoxia alters MSCs’ transcriptional profile, promotes proliferation, increases the production of EVs, and alters the EV proteome, which may enhance the therapeutic efficacy of MSC-EVs ^18,33–36^. MSC-EVs produced under hypoxic conditions have enhanced angiogenic ^37^, osteogenic ^38^, and chondrogenic properties ^39^, as well as improved immunomodulatory and wound-healing effects ^40^. These findings highlight the potential of hypoxic preconditioning as a strategy to enhance the regenerative capacity of MSC-EVs and support their application in regenerative medicine. Importantly, sustained hypoxic culture may produce distinct cellular adaptations compared with short-term hypoxic exposure ^41,42^.

Despite growing evidence that hypoxic conditioning combined with 3D culture can enhance the biological activity of MSCs and their EVs, studies investigating the combined effects of these culture strategies on EV functionality remain limited, and experimental approaches are not standardised ^43–45^. Moreover, the methods used to induce hypoxia, and the selection of 3D culture systems require careful consideration. From a manufacturing perspective, the duration of hypoxic conditioning, adaptation of culture supplements to hypoxic conditions, and the ability to continuously collect EVs during long-term culture may all influence EV yield, composition, and process efficiency. The selection of a 3D culture matrix is also an important manufacturing consideration, as matrix composition can influence MSC phenotype, EV characteristics, and regulatory suitability. Animal-free matrices and culture supplements may facilitate future clinical translation and GMP-compliant production of MSC-EV therapeutics. Furthermore, the proteome of MSC-EVs generated under combined hypoxic and 3D culture conditions has not been comprehensively characterised.

In this study, we evaluated the impact of hypoxia and two animal-free 3D culture platforms on MSC growth, EV production, cargo composition, and regenerative activity. Comprehensive proteomic analyses were performed to characterise EV protein content, and functional activity was assessed using an in vitro scratch wound assay. To our knowledge, this represents the first functional comparison of animal-free hypoxic 3D culture systems for MSC-EV production and provides insights relevant to the development of clinically translatable MSC-EV therapies.

## 1. MATERIALS AND METHODS

### 1.1. 2D Culture

Human adipose-derived stromal cells (MSCs) (#PT-5006, Donors 45450 and 47050, Lonza) were cultured according to manufacturer’s protocol in a T175 flask with MEM α, nucleosides, GlutaMAX Supplement media (#32571028, Thermo Fisher Scientific, Finland) supplemented with filtered 5% human serum (male, blood type AB, #4522, Sigma-Aldrich). The cells were spilt at in a ratio 1:3 when reaching 80% confluence. In the case of EV production, the serum was changed to EV-depleted human serum ^42^. The cells were cultured simultaneously in normoxic (CO_2_ 5%) and hypoxic conditions in a SCI-tive-N hypoxia workstation (Baker Ruskinn) (5% CO_2_, 5% O_2_, 90% N_2_). For 3D culture experiments, cells were used at passages 4 to 6. The cells were tested negative for mycoplasma using MycoAlert Mycoplasma Detection Assay (LT07, Lonza).

### 1.2. 3D culture comparison of culture conditions

To compare the effects of culture conditions on the phenotype of the MSCs and EV production, the cells were cultured in two different 3D models: a matrix-based method with NFCh (GrowDex, UPM Biomedicals, Lappeenranta, Finland) and a matrix-free method using ultra-low attachment (ULA) plates (Nunclon Sphera, Thermo Fisher Scientific). Half of the cells were transferred to hypoxia 4 to 7 days (depending on the growth rate of the cells) before starting 3D culture. Human serum was changed to EV-depleted serum 1 day before beginning 3D culture.

For ULA, 100 μL of cells were seeded at a density of 20 000 cells/mL in ultra-low attachment plates (Nunclon Sphera 96-Well, Thermo Scientific). The cells were allowed to form spheroids undisturbed for 3 days, after which 80 µL/well of medium was refreshed. For NFC, 100 μL of cells were seeded at 500 000 cells/mL in 0.2% (w/v) fibre concentration NFCh in 96-well plates (BRAND microplate BRANDplates, BR781913) as described in ^46^. On day 6, the medium was collected and replaced with fresh medium supplemented with cellulase enzyme (GrowDase, UPM Biomedicals, Lappeenranta, Finland) according to manufacturer’s protocol to digest the hydrogel. The experiment ended on day 7, and the medium was collected for further EV isolation and cells lysed in QIAzol Lysis Reagent (#79306, QIAGEN) and stored at -80 °C.

### 1.3. Viability assessment

The viability of the cells was evaluated by measuring the metabolic activity of the cells at days 2-3 and 6-7 using resazurin reduction assay (alamarBlue, # DAL1025, Invitrogen) according to manufacturer’s protocol: The reagent was added to a final concentration of 10% and incubated for 24 hours. The medium was collected, centrifuged at 150 g for 5 min and the supernatant were collected. The fluorescence intensity was measured from supernatants using a Varioskan LUX multimode microplate reader (Thermo Fisher Scientific); excitation 560 nm, emission 590 nm. The intensities were normalised based on the starting cell amounts.

### 1.4. RNA purification and quantitative PCR

Total RNA was extracted from cell lysates by phenol-chloroform extraction, quantified using a NanoDrop One spectrophotometer (Thermo Fisher Scientific), and reverse-transcribed using the High-Capacity RNA-to-cDNA Kit (#4387406, Applied Biosystems) (Document S1). Relative gene expression was analysed by RT-qPCR using SYBR Green or TaqMan chemistry according to the manufacturer’s instructions. Primer sequences are provided in Table S1, and expression levels were calculated using the 2^-ΔΔCt method^47^.

cDNA from induced pluripotent stem cells was used as external control. The induced pluripotent stem cell line iPS (IMR90)-4, (WiCell, USA) was cultured on Matrigel matrix (#356231, Corning) - coated 6-well plates (#83.3920, Sarstedt, Germany) with 1 mg Matrigel per well, in Essential 8 medium (#A1517001, Thermo Fisher Scientific, USA) in a humified culture incubator (37 °C, 5 % CO2). These cells were previously characterised in ^48^.

### 1.5. Scaling up of 3D culture for EV production for scratch wound experiment and proteomic analysis

Based on the results from the comparative 3D cell experiment, matrix-free ULA model was selected for further studies for upscaled EV production. The cells were seeded at concentration of 1.11 * 10^6^ cells/mL on Corning Elplasia 24-well micropatterned ULA plate (#4441, Corning) according to manufacturer’s protocol, and fresh medium was added on day 3. On day 6 half of the medium was refreshed, and starting on day 8, half of the medium was collected and refreshed every other day until day 14, when all the media were collected. For proteomic analysis the cells were washed with ice-cold 1 × phosphate-buffered saline without calcium or magnesium (-PBS, #70013032, Gibco) and lysed in 8 M urea (#28877.292, VWR Chemicals) in ultrapure water and incubated on ice. The lysate was mixed by vortexing. The samples were stored in -80°C.

### 1.6. Extracellular vesicle purification and characterisation

Conditioned medium was centrifuged at 2500 g for 25 min at 4°C to remove cell debris, and the supernatants were stored at -80°C for further purification. The EVs were purified from the supernatants by differential centrifugation. First the supernatants were centrifuged at 20 000 g for 1 hour to pellet larger EVs and the supernatant was further centrifuged at 110 000 g for 2 h to pellet the remaining small EVs. The pellets were resuspended in either PBS- or 20 mM HEPES (Media Kitchen, University of Helsinki, Finland) and stored -80°C for further analysis. The particle concentration was measured by Nanoparticle Tracking Analysis (Particle Metrix Zetaview PMX-120) performed by EV Core unit at University of Helsinki. The protein content of the EV samples were determined using Pierce BCA Protein Assay Kit (#23227, Thermo Scientific) according to manufacturer’s protocol.

### 1.7. LC-MS total proteome analysis

Proteomic analyses were performed by the Proteomics Unit Viikki, Protein-Proteome Network, University of Helsinki. Detailed procedures regarding total proteome analysis are provided in the Document S1. Shortly, cell and EV lysates were prepared for liquid chromatography-tandem mass spectrometry (LC-MS/MS), and protein concentrations were determined using a BCA assay. Equal amounts of protein (50 µg) from each sample were processed by reduction, alkylation, tryptic digestion, and peptide desalting prior to analysis. Peptide samples were analysed using an Evosep One liquid chromatography system coupled to a timsTOF Pro 2 mass spectrometer (Bruker Daltonics) operating in dia-PASEF mode. Sample-specific acquisition parameters were optimised using data-dependent acquisition runs. Raw data were processed using DIA-NN (v23.0) with a spectral library generated from the UniProt human proteome database^49^. Differentially detected proteins were ranked according to differences in mean spectral counts between culture conditions. Functional enrichment analyses of biological processes were performed using STRINGDB, and protein overlaps between conditions were visualised using UpSet plots generated in R^50^.

### 1.8. *In vitro* Scratch wound assay

To evaluate the therapeutic efficacy and functionality of the EVs, *in vitro* scratch wound assay was performed as described previously ^51^. Briefly, normal human epidermal keratinocytes (NHEKs) (pooled donors, adult, C-12006, Promocell) were cultured according to manufacturer’s protocol in Keratinocyte Growth Medium 2 (C-20011, Promocell) at CO_2_ 5%. A day prior starting the experiment the cells were seeded into Incucyte Imagelock 96-well Microplate (#4379, Sartorius) at density 30 000 cells/well and incubated for overnight to let the cells attach.

The scratch wounds were made using Incucyte 96-well Woundmaker Tool according to manufacturer’s instructions. The EV and platelet lysate (PLTGold Human Platelet Lysate #SCM151, Sigma Aldrich) treatments mix in medium were added to the wells and incubated for 1 to 2 hours before imaging. The imaging was performed using an Incucyte S3 imaging incubator (Software version 2024A). The assay was monitored for 48 or 72 h depending on the wound closure time, and relative wound density and relative wound confluence were subsequently calculated using the built-in analysis Incucyte Scratch Wound Analysis Software Module.

### 1.9. Statistical analysis

Data were tested for normality prior to statistical analysis. Normally distributed data were analysed using one-way ANOVA and Tukey’s HSD Post-hoc Test, as appropriate. Non-normally distributed data were analysed using the Mann–Whitney U test or Kruskal–Wallis test. Statistical analyses and figures were performed using GraphPad Prism version 10.1.2(324), and p < 0.05 was considered statistically significant. The number of replicates (n) refers to biological replicates. Each biological replicate represents the mean of duplicate or triplicate technical replicates.

## 2. RESULTS

### 2.1. Mesenchymal stromal cell phenotype was preserved in long term hypoxia and 3D culture

Two animal-free 3D models were studied: ultra-low attachment (ULA) conditions and nanofibrillated cellulose hydrogel (NFCh). To ensure that MSCs preserved their phenotype, the relative gene expression of selected markers was evaluated after at least 4 days of hypoxic 2D culture and 7 days of hypoxic 3D culture. Cells in all conditions expressed typical MSC markers CD73, CD90, and CD105 (Figure 1A). Stemness markers NANOG, OCT4 and SOX2 were all expressed at lower levels in MSCs than in iPSCs. Paracrine signalling markers VEGF, IL-6 and MCAM did not vary between the cell Matrix and hypoxia impacted EV yield and viability of MSCs

**Figure 1.**
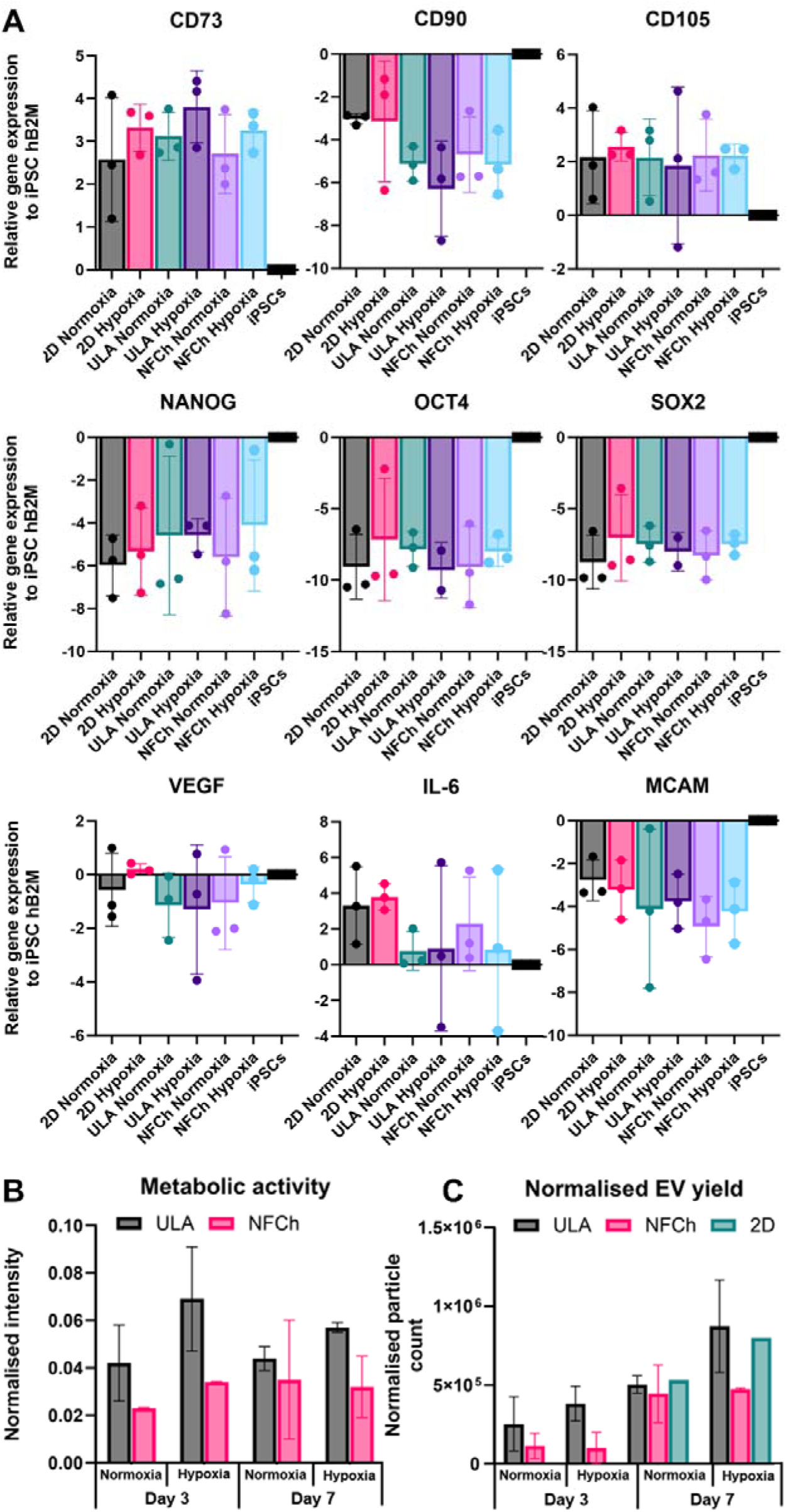
A: MSC gene expression relative to human iPSCs with beta-2-microglobulin calibration. Data are represented as mean ± SD. n=3. There were no statical differences between conditions (p < 0.05). B: Normalised viability of cells and C: normalised EV yield in ULA and NFCh models at day 3 and 7. Normalisation based on seeded cell amounts. Data are mean ± SD (n=2). Abbreviations: ULA = ultra-low attachment plate, NFCh = nanofibrillated cellulose hydrogel. **ALT TEXT :** Multi-panel figure showing mesenchymal stromal cell characterization under normoxic and hypoxic culture conditions. Panel A presents relative gene expression profiles, while panels B and C show normalised cell viability and extracellular vesicle production in ultra-low attachment and nanofibrillated cellulose hydrogel cultures at days 3 and 7.

The viability of cells in different 3D cultures was compared by resazurin reduction (Figure 1B). The ULA model had higher metabolic activity than the NFCh model at day 3 in both normoxia and hypoxia when normalised to starting cell number (Figure 1B). Hypoxia also increased metabolic activity compared to normoxia. On day 7, cells in ULA conditions had the highest viability, but the differences between the models were decreased compared to day 3 (Figure 1B).

The EV yield was greater in ULA than in NFCh at day 3 when normalised to starting cell number (Figure 1C). In addition, hypoxia increased EV yield in ULA but not NFCh. On day 7 ULA and 2D control had similar EV yields in normoxia and hypoxia, both were greater than NFCh. The ULA model was selected for proteomics and *in vitro* scratch wound experiments based on these results and is referred as 3D culture in further experiments. The purity of isolated EVs was evaluated by particles/µg protein ratio, which was consistent with previously reported values ^52,53^ (Table S2). The particle-to-protein ratio was reduced in 3D EV preparations compared with 2D EV preparations, suggesting increased protein content per particle. There was no difference in mean particle size between hypoxic and normoxic EVs but there was a nonsignificant increase in mean particle size 2D EVs compared to 3D EVs (Table S2).

### 2.2. Effect of hypoxia and 3D in proteomic profile of MSCs

LC-MS/MS proteomic analysis identified 7,524 proteins across MSC and EV samples, with MSCs maintaining expression of the characteristic MSC markers CD73, CD90, and CD105 under all culture conditions (Data S2). Proteins commonly expressed across all cellular conditions were primarily associated with gene expression, RNA processing, and peptide metabolism (Data S1). The number of shared proteins across conditions in cells and EVs are presented in Figure 2. For cells, no functional enrichments of Biological processes (BPs) were detected in the highlighted subconditions (Figure 2A, Data S1).

**Figure 2.**
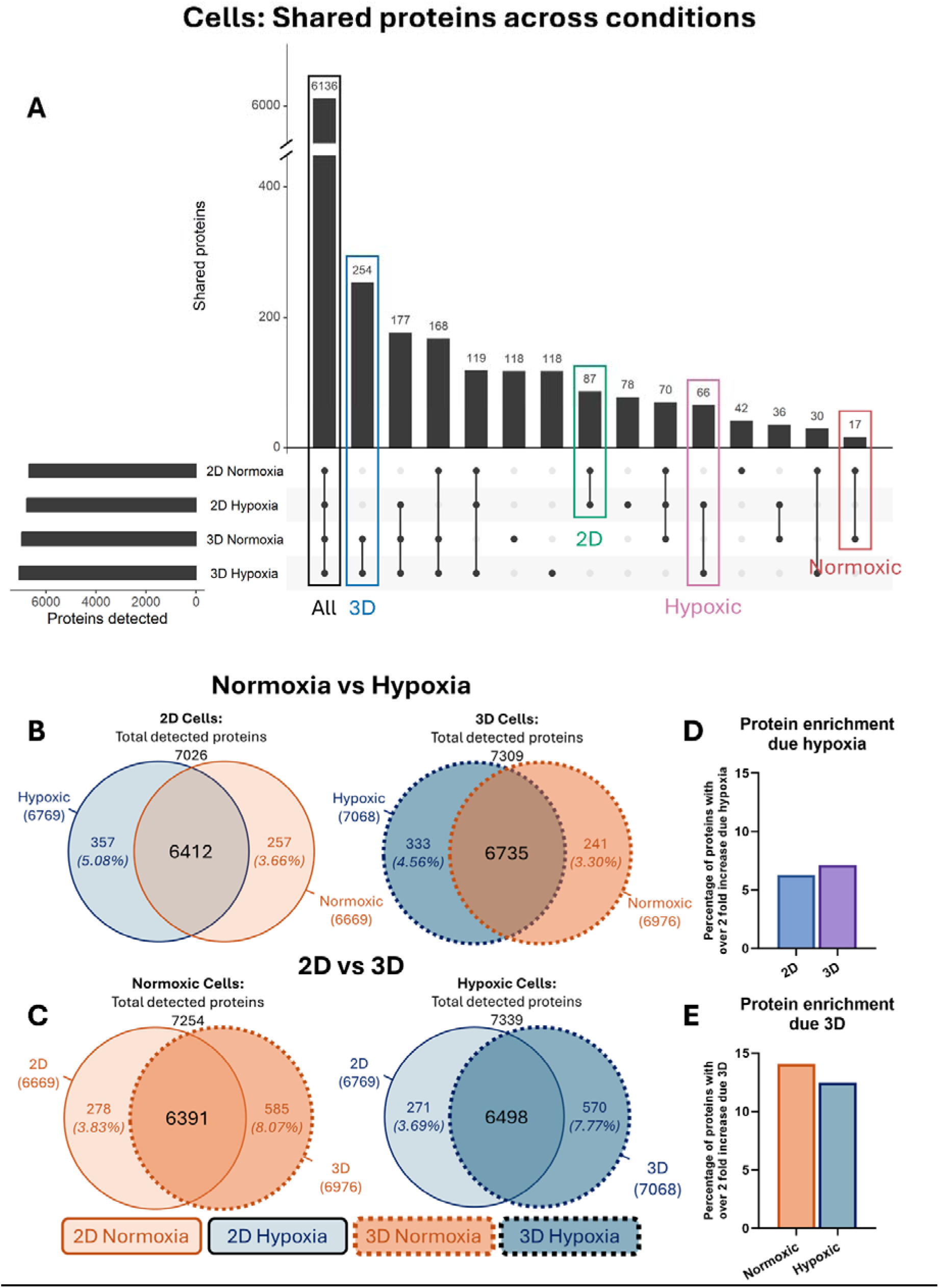
Proteomic overview. Cells: **A:** Number of shared proteins across all conditions. **B:** The impact of hypoxia on shared proteins in 2D and 3D cells. **C:** The impact of 3D culture on shared proteins in normoxic and hypoxic cells. **D:** Proportion of proteins expressed ≥2-fold due to hypoxia in 2D and 3D cells. **E:** Proportion of proteins expressed ≥2-fold due 3D in normoxic and hypoxic cells. **ALT TEXT** : Multi-panel figure showing proteomic comparisons of mesenchymal stromal cells cultured under normoxic and hypoxic two-dimensional and three-dimensional conditions. Panel A shows the overlap of proteins detected across culture conditions. Panels B and C illustrate the effects of hypoxia and three-dimensional culture on protein distribution, respectively. Panels D and E present the proportions of proteins with at least two-fold increased expression associated with hypoxia or three-dimensional culture.

Hypoxia induced relatively modest changes in the cellular proteome, with similar effects observed in both 2D and 3D cultures (Figure 2B, D). Proteins enriched under hypoxic conditions were associated with biological processes related to gene expression, nucleic acid regulation, DNA damage response, cellular stress responses, and metabolism, particularly in 3D cultures (Table 1; Table S3). The 10 most enriched biological processes and proteins due to hypoxia in 3D cells are presented in table 1. A complete list of proteins presented in Data S3.

**Table 1.**
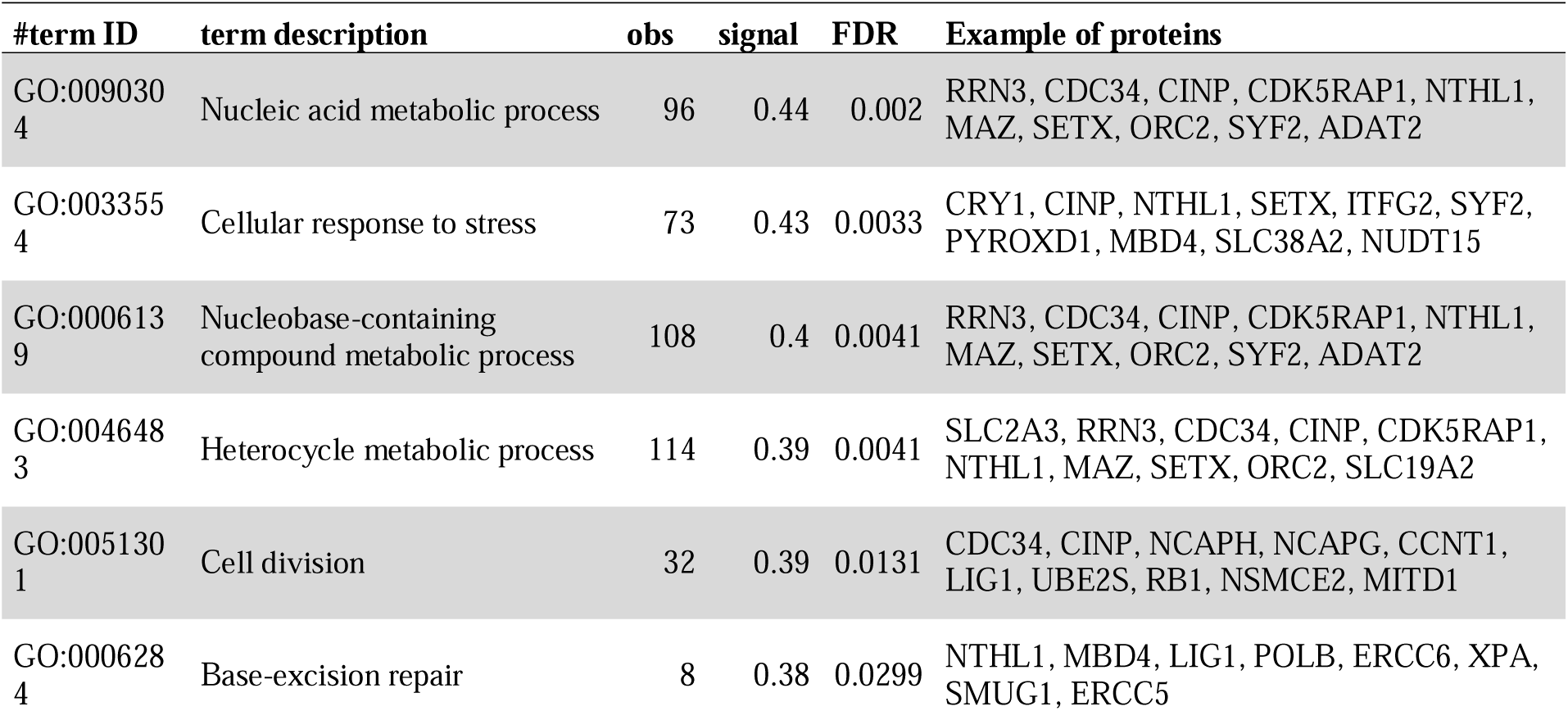
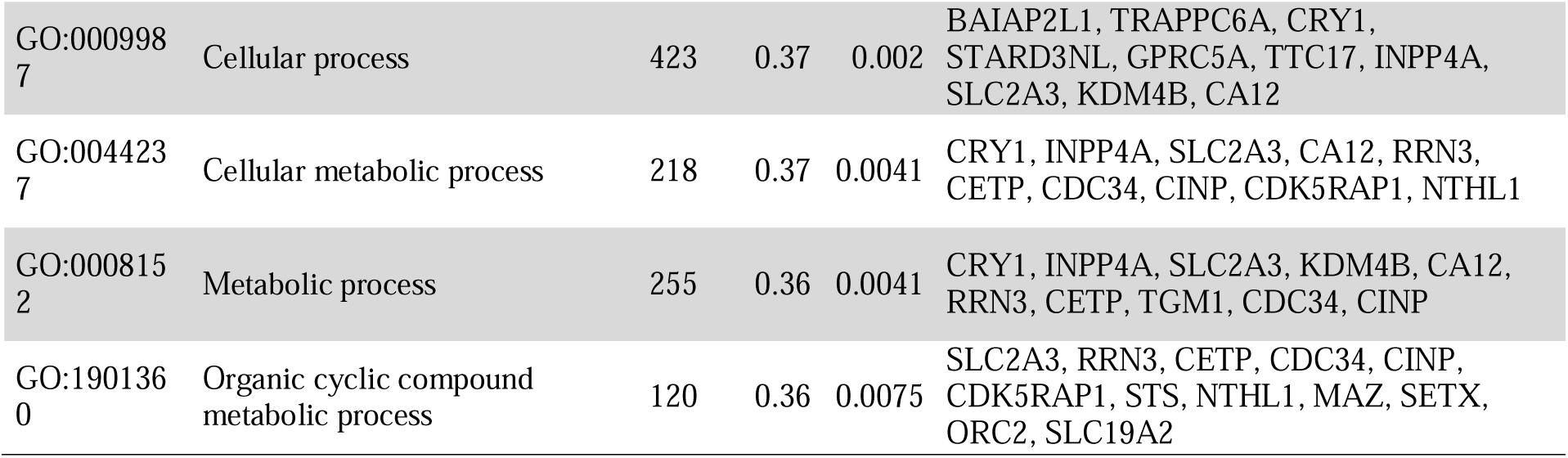
The enriched biological processes and example of enriched proteins due to hypoxia in 3D cells. Completed list of proteins presented in Data S3.

In contrast, 3D culture exerted a greater effect on the MSC proteome than hypoxia alone (Figure 2C, E, Table S4). Proteins enriched under 3D culture conditions were predominantly associated with complement activation, immune-related processes, wound healing, and blood coagulation (Data S3). Comparison of hypoxic 3D cultures with normoxic 2D cultures further highlighted enrichment of proteins involved in DNA processing and repair. Approximately half of the enriched proteins were exclusively detected either in 3D normoxic (585 proteins) or 3D hypoxic (570 proteins) conditions (Table S4). A ranked list of all biological process related to these proteins is presented in Data S3.

Detailed protein identifications, differential expression analyses, functional enrichment results, and complete protein lists are provided in Data S1-S3, and Tables S3-S5.

### 2.3. Effect of hypoxia and 3D in proteomic profile of EVs

LC-MS/MS identified 1,434 proteins across all EV samples, of which 464 were detected under all culture conditions (Figure 3A). Common EV proteins were primarily associated with cytoplasmic translation, humoral immune responses, blood coagulation, and peptide processing (Data S4). EVs from all conditions expressed MSC-associated markers CD73, CD90, and CD105 together with canonical EV marker CD81 and negative for CD63, indicating preservation of EV identity across culture conditions (Data S2). EVs were also positive for the microvesicle markers Calnexin and HSP90B1 ^54^.

**Figure 3:**
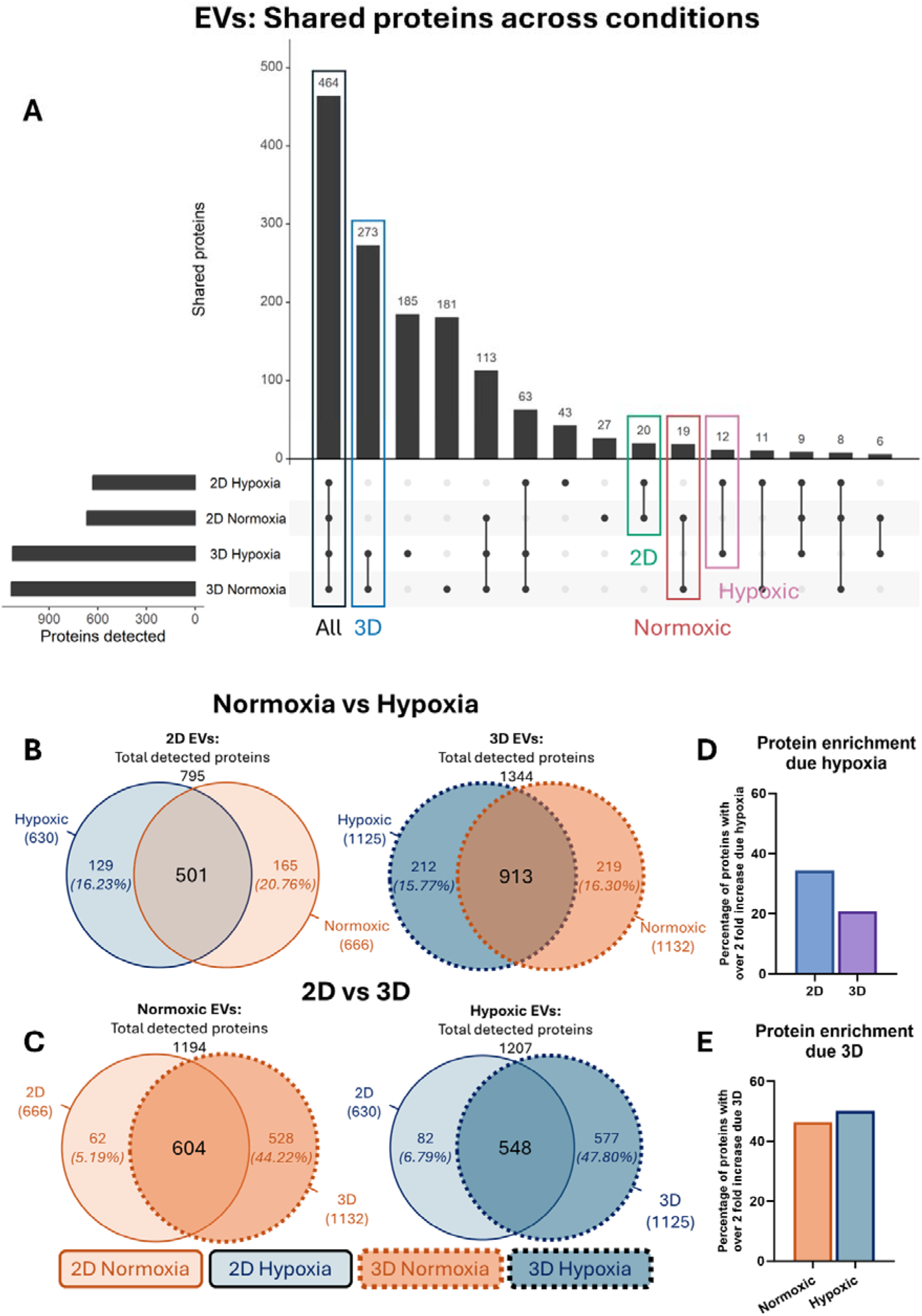
EVs **A:** Number of shared proteins across all conditions. **B:** The impact of hypoxia on shared proteins in 2D and 3D culture derived EVs. **C:** The impact of 3D culture on shared proteins in EVs derived from normoxic and hypoxic cells. **D:** Proportion of proteins expressed ≥2-fold due hypoxia in EVs derived from 2D and 3D cells. **E:** Proportion of proteins expressed ≥2-fold due 3D culture in EVs derived from normoxic and hypoxic cells. **ALT TEXT** : Multi-panel figure showing proteomic comparisons of extracellular vesicles derived from mesenchymal stromal cells cultured under normoxic and hypoxic two-dimensional and three-dimensional conditions. Panel A shows the overlap of proteins detected across extracellular vesicle preparations. Panels B and C illustrate the effects of hypoxia and three-dimensional culture on extracellular vesicle protein composition, respectively. Panels D and E present the proportions of proteins with at least two-fold increased expression associated with hypoxia or three-dimensional culture in extracellular vesicles.

Hypoxia altered EV protein cargo in both 2D and 3D cultures (Figure 3B, D; Table S6). Total number of detected proteins was higher in EVs derived from 3D culture (1344) compared to 2D (795). In 2D hypoxic EVs, 34.3% of proteins had ≥2-fold increase, compared to 20.833% in hypoxic 3D EVs (Figure 3D). In 2D cultures, proteins enriched under hypoxia were primarily associated with cell adhesion, cell signalling, and haemostasis, whereas proteins enriched in hypoxic 3D EVs were predominantly related to translation and peptide-processing pathways (Data S4).

Compared with hypoxia, 3D culture exerted a substantially greater effect on EV composition (Figure 3C, E; Table S7). A notable increase in proteins with ≥2-fold increase in expression was observed in 3D EVs, accounting for 46.3% in normoxia and 50.1% in hypoxia (Figure 3E). This was due to a substantial fraction of proteins uniquely detected in 3D EVs (528 proteins, 44.2% in normoxia; 577 proteins, 47.8% in hypoxia). EVs derived from 3D cultures contained a larger number of detectable proteins and showed extensive enrichment of proteins involved in peptide processing and protein metabolism under both oxygen conditions (Data S4). Enrichment of proteins with ≥2-fold increase in expression in 3D EVs across conditions and the top 10 biological processes and example of proteins are presented in Table 2 and Data S4.

**Table 2.** The ten most enriched biological process based on signal in 3D EVs compared to 2D EVs and example of proteins. For complete list of proteins see Data S4.

| #term ID | Biological process | obs. | signal | FDR | Example of proteins |
|---|---|---|---|---|---|
| GO:0002181 | Cytoplasmic translation | 77 | 4.16 | 1.26E-37 | EIF3D, RPL18A, RPL19, RPS12, EIF2S3, RPL35, RPL8, RPS11, RPS27A, RPS3 |
| GO:0006956 | Complement activation | 41 | 3.21 | 4.06E-21 | C5, C1QBP, C4BPB, C3, CFP, KRT1, CFHR5, C9, C6, C1RL |
| GO:0006958 | Complement activation, classical pathway | 29 | 2.66 | 2.07E-15 | C5, C1QBP, C4BPB, C3, C9, C6, C1RL, SERPING1, C2, C1QB |
| GO:0002455 | Humoral immune response mediated by circulating immunoglobulin | 30 | 2.61 | 1.97E-15 | C5, C1QBP, C4BPB, C3, C9, C6, CD81, C1RL, SERPING1, C2 |
| GO:0006412 | Translation | 119 | 2.55 | 3.97E-35 | EIF3D, EEF1A2, RPL18A, RPL19, RPS12, RARS1, EIF2S3, EIF2S1, NARS1, RPL35 |
| GO:0043043 | Peptide biosynthetic process | 121 | 2.43 | 5.58E-34 | EIF3D, EEF1A2, RPL18A, RPL19, RPS12, RARS1, EIF2S3, EIF2S1, NARS1, RPL35 |
| GO:0007596 | Blood coagulation | 64 | 2.43 | 5.29E-22 | CPB2, APOH, VCL, SERPIND1, MYH9, PROCR, F9, COMP, TFPI2, HRG |
| GO:0007599 | Haemostasis | 65 | 2.42 | 4.24E-22 | CPB2, APOH, VCL, SERPIND1, MYH9, PROCR, F9, COMP, TFPI2, HRG |
| GO:0006518 | Peptide metabolic process | 144 | 2.31 | 8.96E-37 | EIF3D, EEF1A2, RPL18A, RPL19, LTA4H, RPS12, RARS1, EIF2S3, EIF2S1, NARS1 |
| GO:0042060 | Wound healing | 97 | 2.25 | 8.32E-27 | CPB2, APOH, VCL, SERPIND1, MYH9, PROCR, F9, TIMP1, TGFB1, COMP |

Overall, while both hypoxia and 3D culture modified EV cargo composition, 3D culture produced the most pronounced changes, indicating that culture dimensionality is a major determinant of MSC-EV protein content.

### 2.4. EV vs cell proteomics

EV proteomes were compared to the parental cells to assess protein enrichment in EVs relative to the cellular proteome of host cells. EVs had notably fewer proteins exclusively expressed (<1%) compared to cells in all conditions (Table S8 & S9). Despite this, approximately 4-5% (507 proteins) of detected proteins in EVs had ≥2-fold increase in expression compared to cells. Of the 507 enriched proteins in EVs, 204 were common across all conditions and were identified by enrichment analysis as serum-associated proteins (Figure 4, Data S5).

**Figure 4.**
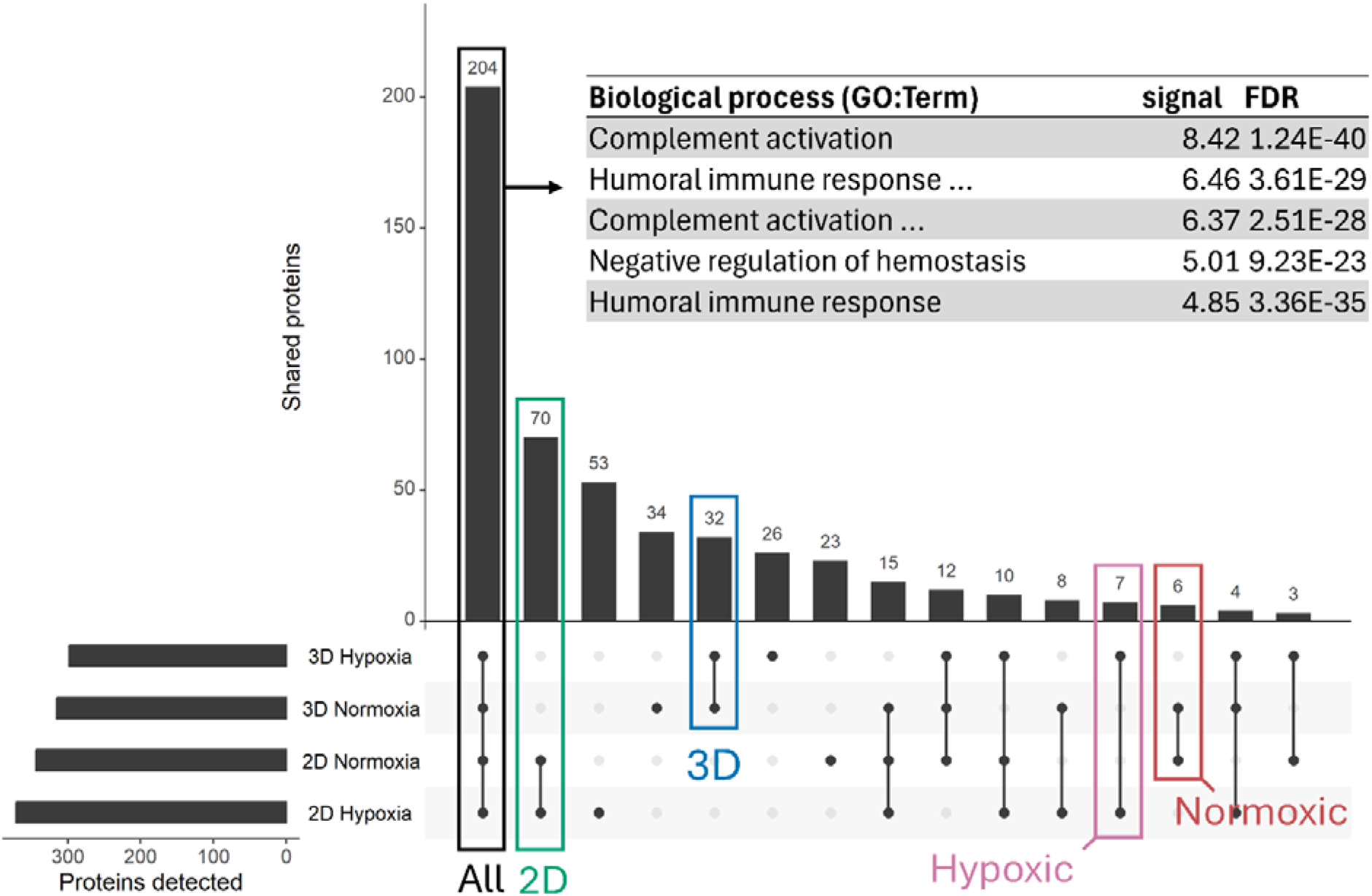
Number of shared proteins across all conditions with ≥2-fold enrichment in EVs compered to their host cell. Total amount of proteins with ≥2-fold enrichment in EVs compered to their host cell = 507. Complete list of proteins and enriched biological processes in presented in Data S5. **ALT TEXT :** Figure showing proteins enriched in extracellular vesicles relative to their parent mesenchymal stromal cells across all culture conditions. The diagram displays the overlap of proteins with at least two-fold higher abundance in extracellular vesicles than in the corresponding host cells and highlights proteins shared between experimental groups. Detailed protein identifications and associated biological processes are provided in the supplementary data.

### 2.5. Hypoxia improved the wound healing efficacy of 3D EVs *in vitro*

The impact of 3D culture and hypoxia on the therapeutic efficacy of MSC-EVs was studied using an *in vitro* scratch wound assay and measuring gene expression of genes of interest in normal human keratinocytes (NHEKS) after EV treatment (Figure 5). Wounds treated with 2D EVs closed the fastest regardless of oxygen condition (80% closure in 10 hours), faster than the 5% platelet lysate positive control (80% closure in 12 hours) (Figure 5B, Table S10). Hypoxia improved the therapeutic efficacy of 3D EVs; however, the effect (80% closure in 18 hours) was less than 2D EVs. In both the normoxic 3D EV-treated group and the negative control, 80% wound closure took longer than 24 hours and complete closure did not occur within 48 hours, unlike other experimental conditions.

**Figure 5.**
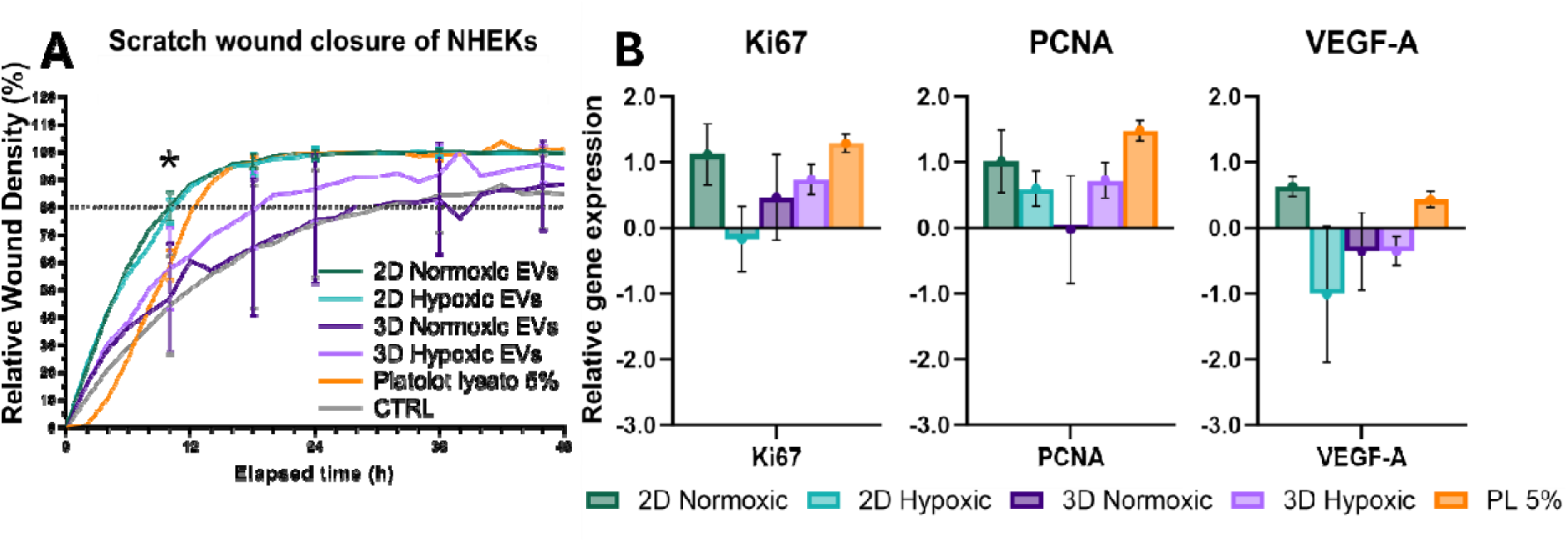
Wound healing efficacy of MSC-EVs from hypoxic and 3D culture conditions in a 48-hour NHEK scratch wound model. **A:** The impact of EVs on the closure time measured as relative wound density. * = 2D normoxic and hypoxic EVs vs CTRL (p < 0.05) using one-way ANOVA and Tukey’s HSD Post-hoc Test (n=4). Statical analysis of the selected timepoint presented in Figure S1 and Table 10. **B:** Fold change in relative gene expression of Ki67, PCNA and VEGF-A in NHEKs after a 48-hour scratch wound assay treated with MSC-EVs from hypoxic and 3D culture conditions compared to relative to untreated GAPDH calibration. **ALT TEXT** : Multi-panel figure showing the effects of extracellular vesicles derived from mesenchymal stromal cells cultured under normoxic, hypoxic, and three-dimensional conditions in an in vitro keratinocyte scratch wound model. Panel A presents wound closure over 48 hours measured as relative wound density across treatment groups. Panel B shows relative expression of the proliferation markers Ki67 and PCNA and the angiogenesis-related gene VEGF-A in keratinocytes following extracellular vesicle treatment.

After a 48-hour scratch wound assay, the expression of the proliferation markers Ki67 and PCNA, as well as VEGF-A, were measured relative to untreated controls (Figure 5B). Ki67 was non-significantly upregulated in all conditions except 2D hypoxic EVs. PCNA was non-significantly upregulated in all conditions except 3D normoxic EVs. VEGF-A was upregulated in 2D normoxic EVs and 5% platelet lysate and downregulated in other conditions. These results aligned with those observed at 72 hours (Figure S2).

## 3. Discussion

In this study, the phenotype of MSC-EVs derived from two different animal-free 3D models in hypoxic conditions was compared. The proteome of these EVs and their parent cells were analysed in addition to relevant gene expression.

Previous studies have shown that hypoxia and 3D culture can individually enhance the therapeutic potential of MSCs and MSC-derived EVs; however, their combined effects remain poorly characterised, particularly under long-term hypoxic conditions ^43,44,55,56^. Most studies employ short-term hypoxic exposures that primarily capture acute cellular responses, whereas sustained hypoxic culture may induce distinct adaptive changes relevant to continuous EV production ^34,41,57–59^. The reported effects of hypoxia on MSC proliferation, viability, and stemness have also been variable, likely reflecting differences in MSC source, donor heterogeneity, and culture conditions ^34,60–62^.

From a translational perspective, both hypoxia and 3D culture represent potentially important manufacturing parameters that can influence EV yield, composition, and therapeutic activity ^55,56,63,64^. In the present study, long-term hypoxia increased EV production in matrix-free 3D culture, supporting its potential use as a process optimisation strategy for MSC-EV manufacturing. In contrast, the hydrogel-based culture system retained EVs within the matrix and required hydrogel digestion for recovery, limiting its suitability for scalable EV production^65^. These findings highlight the importance of considering not only EV bioactivity but also product recovery and manufacturability when developing clinically translatable MSC-EV production platforms.

### Hypoxia and 3D culture elicit similar proteomic responses in MSCs

The MSC markers CD73, CD90 and CD105 were detected in both qPCR and proteomics, and the negative markers CD11b, CD45, CD34, CD19, CD79a, and HLA-DR were undetected supporting

MSC identity according to the ISCT criteria ^1^. In contrast to several reports describing increased stemness following hypoxic preconditioning or 3D culture, neither hypoxia nor 3D culture increased expression of NANOG, SOX2, or OCT4 in the present study ^20–24,60–62,66^. This finding agrees with recent studies that prolonged hypoxic culture primarily promotes cellular adaptation rather than transient stress responses associated with enhanced stemness ^60–62^. We suggest that long-term hypoxic conditioning promotes cellular adaptation rather than stress-induced responses, explaining the absence of increased stemness, or paracrine signalling compared with reports using short-term hypoxic preconditioning.

Proteomic analyses further supported successful adaptation to long-term hypoxia, with enrichment of proteins associated with cellular metabolism, stress responses, and genome maintenance. Notably, 3D culture had a greater impact on the MSC proteome than oxygen tension alone, suggesting that culture architecture may be a major determinant of MSC phenotype. The more limited response to additional hypoxic stimulation observed in 3D cultures may reflect partial adaptation to reduced oxygen availability within spheroid structures. Lack of HIF-1α, a regulator of hypoxic adaptation, in hypoxic cells is likely since HIF-1α regulates the acute hypoxic response and the MSCs are already adapted to hypoxia ^67–69^. Enrichment of hypoxia associated proteins AJUBA and GLUT in hypoxic cells may indicate hypoxia-induced metabolic reprogramming toward glycolysis and hypoxia-associated remodelling of adhesion and HIF-related signalling pathways ^70,71^. Interestingly, CA9 and STC2 both regulated by the HIF pathway, were detected exclusively in 3D hypoxic cells. This supports our assertion that 3D cultures are inherently more hypoxic than 2D cultures ^72,73^. Cells in 3D culture form a solid spheroid limiting oxygen access to the core ^74^. Therefore, 3D cultures may be partially preconditioned to a hypoxia-adapted state, even under normoxic conditions, resulting in a less coordinated proteomic response to additional hypoxia.

From a translational perspective, these findings indicate that prolonged hypoxia and 3D culture do not enhance conventional MSC stemness markers but instead induce broader changes in cellular state and protein expression. Such changes may influence EV composition and function and therefore represent important manufacturing variables during development of MSC-EV therapeutics. When considering only proteins that were upregulated ≥2-fold in hypoxia, 3D cells were significantly enriched with proteins associated with cellular stress and cellular metabolism, metabolism of nucleic acids, genome maintenance and stress response, and cellular growth and proliferation indicating adaptation to a changing culture environment. These adaptations could affect the regenerative capacity of MSCs, and if also present in EVs, could be important for MSC-EV therapies. It is remarkable that similar biological processes are enriched in 2D and 3D given that an improvement in wound healing capacity was only observed in 3D EVs.

Recent evidence has challenged the concept that hypoxia or 3D culture enhance MSC stemness and proliferation ^20–24,62,66^. *In vivo*, MSCs are largely quiescent and are activated only in response to external cues. Therefore, hypoxic 3D culture systems that resemble the *in vivo* niche may maintain MSCs in a dormant state. Further studies should evaluate the functional potential of MSCs under these conditions.

Human serum was used to support MSC growth in this animal-free culture system and contributed serum-associated proteins to both cellular and EV proteomes. Because serum supplementation was identical across all conditions, comparative analyses focused on proteins uniquely detected or enriched in specific culture conditions. Increased abundance of serum-associated proteins in 3D cultures may reflect culture-dependent alterations in protein association and could influence EV biodistribution or uptake, as suggested previously ^75–77^.

### 3D EVs are more protein rich but perform more poorly in wound healing than 2D EVs

EVs from all culture conditions retained expression of MSC-associated markers, confirming their MSC origin^78^. Calnexin was also detected, which is commonly observed in EV preparations isolated by ultracentrifugation^54^.

Proteomic analyses revealed increased protein abundance complexity in EVs derived from 3D cultures compared with 2D cultures under both normoxic and hypoxic conditions. Although serum-associated co-isolated proteins were detected in all EV preparations^52^, their relative enrichment differed between culture conditions, suggesting that culture configuration influences the protein composition associated with EVs. Previous studies have shown that 3D culture-dependent changes in the EV protein corona can affect cellular uptake, biodistribution, and biological activity supporting our findings ^76,79^.These findings highlight the importance of culture conditions in determining EV function ^75,77,80^.

Despite their greater proteomic complexity, 3D-derived EVs did not improve wound healing compared with 2D-derived EVs. Furthermore, comparison of hypoxic 3D EVs with normoxic 2D EVs suggested that 3D culture exerted a larger influence on EV protein composition than hypoxic conditioning alone. Together, these findings indicate that increased EV protein content does not necessarily translate into improved therapeutic efficacy. From a translational perspective, functional potency assays remain essential for evaluating MSC-EV products, as molecular complexity alone may not predict biological performance.

When EV protein content was compared with that of the parent MSCs, proteins associated with serum-derived biological processes were enriched in all EV preparations (Figure 4 (Data S5)). Because EVs were isolated by ultracentrifugation which is sensitive for affecting the EV corona, it was not possible to distinguish between co-isolated soluble proteins and proteins associated with an acquired EV corona ^81,82^. Since the protein corona and other EV-associated proteins play crucial roles in cellular uptake and biodistribution both *in vitro* and *in vivo*, their contribution to the therapeutic efficacy of EVs should not be overlooked. Future studies should therefore consider EV corona composition as a potentially important quality attribute during MSC-EV manufacturing and characterization.

### Effect of culture conditions on the therapeutic potential of EVs

The therapeutic effects of MSC-EVs have been demonstrated in numerous *in vitro* and *in vivo* wound-healing models, but the combined impact of hypoxia and 3D culture on EV functionality remains poorly understood.^83–85^. In the present study, EVs derived from 2D cultures promoted scratch wound healing more effectively than EVs derived from 3D cultures or platelet lysate, irrespective of oxygen condition. Normoxic 2D EVs and platelet lysate had similar effects increasing the gene expression of Ki67, PCNA and VEGF-A in NHEKs. Hypoxic 3D EVs also increased Ki67 and PCNA expression, but surprisingly downregulated VEGF-A. Although hypoxic MSCs are frequently associated with enhanced pro-angiogenic activity, this effect may not necessarily be mediated through increased VEGF expression. Furthermore, keratinocytes are not the primary regulators of angiogenesis, and reduced VEGF expression in NHEKs does not necessarily reflect diminished angiogenic potential.

However, previous reports showed similar improvement in proliferation, migration and wound healing in human keratinocytes treated with MSC normoxic 2D EVs ^86,87^. Yuan et al. demonstrated that hypoxic preconditioning increased the expression of Ki-67 in hypoxic renal injury in rats ^36^ and others have reported enhanced regenerative capacity of hypoxic preconditioned MSC-EVs especially in the context of wound healing *in vitro* and *in vivo* ^40,88,89^.

Although 3D culture increased protein complexity in both MSCs and their EVs, this did not translate into enhanced therapeutic efficacy in the *in vitro* scratch wound assay. However, the *in vitro* wound model does not fully recapitulate the complexity of the physiological wound healing process, including immune cell interactions, angiogenesis, and tissue remodelling. Therefore, mechanistic *in vivo* wound healing studies are required to determine whether the proteomic changes induced by 3D culture confer additional therapeutic benefits. Nevertheless, EVs from 2D cultures improved wound closure *in vitro* reenforcing the role of MSC-EVs in regenerative applications.

## 4. Conclusion

To our knowledge, this is the first study to evaluate the effects of long-term hypoxia and animal-free 3D culture on the proteomic composition of MSC-derived EVs and to relate these changes to functional wound-healing activity. Long-term hypoxia and 3D culture altered EV production and cargo composition without compromising MSC identity. While 3D culture had the greatest effect on both cellular and EV proteomes, hypoxic conditioning further enhanced EV yield and improved the wound-healing activity of 3D-derived EVs.

Importantly, our findings demonstrate that culture configuration, oxygen tension, and matrix selection are not merely experimental variables but manufacturing parameters that influence MSC-EV yield, composition, recovery, and biological activity. The matrix-free hypoxic 3D culture system supported continuous EV collection and generated EVs with distinct molecular and functional characteristics, highlighting its potential as a clinically relevant production platform.

Although increased proteomic complexity did not directly translate into superior wound-healing efficacy, hypoxia-induced alterations in EV cargo were associated with improved functional performance, emphasizing the importance of combining molecular characterization with biological potency testing during MSC-EV development. Future *in vivo* studies are required to determine how these culture-induced changes influence therapeutic efficacy, biodistribution, and safety. Overall, this work supports the integration of physiologically relevant culture conditions into MSC-EV manufacturing strategies and provides new insights for the development of scalable and clinically translatable MSC-EV therapeutics.

## Author contributions

Julia Monola: Conception and design, Collection and/or assembly of data, Data analysis and interpretation, Manuscript writing, Final approval of manuscript.

Alisa Ottila: Conception and design, Collection and/or assembly of data, Data analysis and interpretation, Manuscript writing, Final approval of manuscript.

Nikolas Heikkilä: Collection and/or assembly of data, Final approval of manuscript.

Chris S. Pridgeon: Conception and design, Data analysis and interpretation, Manuscript writing, Final approval of manuscript.

Riina Harjumäki: Conception and design, Financial support, Administrative support, Provision of study material or patients, Data analysis and interpretation, Manuscript writing, Final approval of manuscript.

## 6. Acknowledgments

Funding was provided by the Research Council of Finland (grant no. 375887 and 337430), the Doctoral Programme in Materials Research and Nanoscience (JM), the Doctoral Programme in Drug Research (AO), and the Finnish Cultural Foundation (grant no. 00250651; JM, 00250755; CP and 00220283; RH).

The authors gratefully acknowledge the following University of Helsinki services: EV core, for NTA services; Proteomics Unit Viikki, Protein-Proteome Network Helsinki for LC-MS proteomic analysis services; the DDCB Unit at the Faculty of Pharmacy, supported by HiLIFE and Biocenter Finland; and the HiLIFE Biocomplex unit, a member of Instruct-ERIC Centre Finland, FINStruct, and Biocenter Finland. The graphical abstract was created using BioRender.

## 7. Data Availability Statement

The mass spectrometry proteomics data have been deposited to the ProteomeXchange Consortium via the PRIDE partner repository with the dataset identifier PXD081617 ^90^. All other data supporting the findings of this study are available from the corresponding author upon reasonable request. This study did not generate custom code.

## 8. Declaration of generative AI and AI-assisted technologies in the manuscript preparation process

During the preparation of this work, the authors used ChatGPT (OpenAI) for grammar and language editing. The authors reviewed and edited the output as needed and take full responsibility for the content of the published article.

