## Supplementary material for "Physiological Culture Modulates the Cargo and Wound-Healing Potential of MSC Extracellular Vesicles": Document S1

### **MATERIALS AND METHODS**

#### RNA purification and quantitative PCR

RNA was purified from the cell lysates by phenol-chloroform RNA extraction. Chloroform was added to the samples in a 1:5 ratio and shaken vigorously for 15 seconds which after the samples were centrifuged at 16,000 × g for 15 minutes at 4°C. The aqueous phase was collected and mixed with an equal volume of isopropanol. The RNA was pelleted by centrifugation at 16,000 × g for 10 minutes at 4°C and washed with 75% ethanol. The ethanol was removed by centrifugation, and the pellet was resuspended in RNA-free water. The RNA yield was measured using NanoDrop One UV-Vis spectrophotometers (Thermo Fisher Scientific). Purified RNA was reverse-transcribed using the High-Capacity RNA-to-cDNA Kit (#4387406, Applied Biosystems) and the cDNA was stored at -20°.

The relative gene expression was studied using real-time quantitative PCR (RT-qPCR). Primers and master mix were used according to manufacturer’s protocols (PowerUP SYBR Green Master Mix A25742, Applied Biosystems) and TaqMan Universal PCR Master Mix No AmpErase UNG (#4324018, Applied Biosystems). The primers used are presented in Table M1. RT-qPCR run parameters were performed according to manufacturer’s instructions. Data analysis was performed using the 2-ΔΔCt method [87].

cDNA from induced pluripotent stem cells was used as external control. The induced pluripotent stem cell line iPS(IMR90)-4, (WiCell, USA) was cultured on Matrigel matrix (#356231, Corning) -coated 6-well plates (#83.3920, Sarstedt, Germany) with 1 mg Matrigel per well, in Essential 8 medium (#A1517001, Thermo Fisher Scientific, USA) in a humified culture incubator (37 °C, 5 % CO2). These cells were previously characterised in [88].

**Table S1.** Details of the used primers used in RT-qPCR assays.

| Cell line | Target gene | F- primer (sequence 5′-3′) | R-primer (sequence 5′-3′) |
| --- | --- | --- | --- |
| MSCs | β-2-M | CTCGCGCTACTCTCTCTTTCTG | GCTTACATGTCTCGATCCCACT |
|  | CD73 | CTTAACGTGGGAGTGGAACC | TCTAGCTGCCATTTGCACAC |
|  | CD90 | CGCTCTCCTGCTAACAGTCTT | CAGGCTGAACTCGTACTGGA |
|  | CD105 | CTAACTGGCAGGGGAGACAG | CTCCATGTGGCAGGAGCTA |
|  | NANOG | GCAGAAGGCCTCAGCACCTA | GGTTCCCAGTCGGGTTCAC |
|  | OCT4 | CAGTGCCCGAAACCCACAC | GGAGACCCAGCAGCCTCAAA |
|  | SOX2 | CTCCGGGACATGATCAGC | GGTAGTGCTGGGACATGTGAA |
|  | IL-6 | AACCTGAACCTTCCAAAGATGG | TCTGGCTTGTTCCTCACTACT |
|  | MCAM | ATCGCTGCTGAGTGAACCACAG | CTACTCTCTGCCTCACAGGTCA |
|  | VEGF | TGCTTCTGAGTTGCCCAGGA | TGGTTTCAATGGTGTGAGGACATAG |
| Cell line | Gene | Primer ID | |
| NHEKs | GADPH | Hs02786624_g1 | |
|  | PCNA | Hs00427214_g1 | |
|  | Ki67 | Hs04260396_g1 | |
|  | VEGF-A | Hs0090055_m1 | |

#### LC-MS total proteome analysis

#### 1. Sample preparation

To compare the effects of culture conditions to the phenotype of the cells and their EVs, LC-MS total proteome analysis was performed by the Proteomics Unit Viikki, Protein-Proteome Network Helsinki University of Helsinki. First, the cell samples were lysed in 8 M urea, and the protein concentrations were determined using BCA assay according to manufacturer’s instruction. The EVs were first lysed in lysis buffer (Tris-Hl, NaCL, Triton X-100) by incubating in ice for 30 min and then centrifuging the samples at 17 000 g for 20 minutes. The protein concentrations of the supernatant was determined using a BCA assay.

50 µg of protein in PBS- from each sample was used for proteomics analysis. 8 M urea (Fisher, cat. ##424585000) in 100 mM AMBIC (Honeywell Fluka, cat. #15661030) was added to the samples, brief vortexed and sonicated for 10 minutes in a water bath. Then the samples were reduced with 5 mM Tris(2-carboxyethyl)phosphine hydrochloride (#20490, Thermo Scientific), alkylated with 10 mM iodoacetamide (#122271000, Acros Organics) at room temperature, and the urea concentration was diluted to 1 M with 100 mM AMBIC. Reduced and alkylated peptides were trypsin-digested at 37°C for 16 hours using Sequencing Grade Modified Trypsin (V5113, Promega). After digestion, samples were acidified with 10% trifluoroacetic acid (Fisher, cat. #10723857) and desalted with BioPureSPN PROTO 300 C18 Mini columns (#HUM S18V, Higgins analytical, USA) according to manufacturer’s instructions. After desalting the samples were dried in a centrifuge concentrator (Concentrator Plus, Eppendorf). The dried peptides were reconstituted in 30 µL buffer A (0.1% TFA, 1% acetonitrile (Fisher, cat. 10001334) in HPLC grade water (Fisher, cat. 10777404)).

For the DIA analysis of samples, the resuspended peptides were further diluted to 7.5 ng/µL in buffer A1 (1% formic acid (#A117-50, Fisher) in HPLC water). 20 µL was loaded into an Evotip (Evosep, Denmark) following manufacturer’s instructions to achieve ~150 ng of peptide load for the analysis.

## 2. LC-MS/MS

The desalted samples were analysed using the Evosep One liquid chromatography system coupled to a hybrid trapped ion mobility quadrupole TOF mass spectrometer (Bruker timsTOF Pro 2, Bruker Daltonics) (Meier, Brunner et al., 2018) via a CaptiveSpray nano-electrospray ion source (Bruker Daltonics). An 8 cm × 150 µm column with 1.5 µm C18 beads (EV1109, Evosep) was used for peptide separation with the 60 samples per day methods (21 min gradient time). Mobile phases A and B were 0.1 % formic acid in water and 0.1 % formic acid in acetonitrile, respectively. The MS analysis was performed in the positive-ion mode with dia-PASEF method [89] with sample optimised data independent analysis (dia) scan parameters. We performed DDA in PASEF mode from a pooled sample to be able to adjust dia-PASEF parameters optimally. To perform sample specific dia-PASEF parameter adjustment the default dia-short-gradient acquisition methods was adjusted based on the sample specific DDA-PASEF run with the software “tims Control” (Bruker Daltonics). The following parameters were modified for each sample type: m/z range: 410.0 – 1072.0 Da; mobility range 0.72 – 1.33; mass window width 25.0 Da with 0.5 Da overlap; mass steps per cycle: 27; mean cycle time: 0.95 s. The ion mobility windows were set to best match the ion cloud density from the sample type specific DDA-runs.

#### **Data processing**:

To analyse diaPASEF data, the raw data (.d) were processed with DIA-NN v23.0 [90,91] using a spectral library generated from the UniProt human proteome (USP000000589, downloaded 06.02.2025 as a FASTA file, 20394 proteins). During library generation following settings were used, fixed modifications: carbamidomethyl (C); variable modifications: acetyl (protein N-term), oxidation (M); enzyme:Trypsin/P; maximun missed cleavages: 1; mass accuracy fixed to 1.5e-05 (MS2) and 1.5e-05 (MS1); Fragment m/z set to 200-1800; peptide length set to 7-50; precursor m/z set to 300-1800; Precursor changes set to 2-4; protein inference not performed. All other settings were left as default.

The difference in mean spectral count between conditions was used to create a ranked list of detected proteins. The “Proteins with Values/Ranks” tool in StringDB version 12.0 was used to generate a list of biological process (BP) GO terms using the Homo sapiens database [92]. Enriched biological processes were identified from proteins that were exclusively detected or showed a ≥2-fold increase in expression in the studied conditions. The enrichment were ranked against the signal. The signal is defined as a weighted harmonic mean between the observed/expected ratio and -log(FDR). Protein intersections between the studied conditions were assessed using UpSet plots generated using R (version 4.6.0) in RStudio (version 2026.01.2) with the UpSetR package [93].

### **RESULTS**

Table S2. EV characteristics: Mean and median particle size (nm) and particles/protein µn to evaluate purity of samples.

| **Cell culture type** | | **Size nm** | | | **Particles/Protein µg** | |
| --- | --- | --- | --- | --- | --- | --- |
|  |  | **Mean** | **Median** | **SD (mean/median)** | **Mean** | **SD** |
| **2D** | Normoxia (n =4) | 134.3 | 119.4 | 3.12 / 1.95 | 1.30E+09 | 2.84E+08 |
|  | Hypoxia (n =4) | 137.6 | 123.2 | 6.17 / 8.21 | 1.77E+09 | 4.83E+08 |
| **3D** | Normoxia (n =3) | 125.8 | 111.8 | 4.74 / 4.74 | 8.30E+07 | 2.16E+07 |
|  | Hypoxia (n =3) | 128.1 | 112.2 | 5.46 / 5.24 | 6.91E+07 | 3.92E+06 |

Table S3. The impact of hypoxia in 2D and 3D cells on the number of detected proteins and the abundance of proteins enrich over 2-fold. Abbreviations: H = hypoxia, N = normoxia.

| **Hypoxia: Cells** | | | | | | | | | |
| --- | --- | --- | --- | --- | --- | --- | --- | --- | --- |
| **Cell culture** | **Total** | **Normoxic** | **Only normoxic** | **%** | **Hypoxic** | **Only hypoxic** | **%** | **H vs N enrichment (2-fold)** | **%** |
| **2D** | 7026 | 6669 | 257 | 3,658 % | 6769 | 357 | 5,081 % | 442 | 6,291 % |
| **3D** | 7309 | 6976 | 241 | 3,297 % | 7068 | 333 | 4,556 % | 522 | 7,142 % |

Table S4. The impact of 3D cell culture in normoxic and hypoxic cells on the number of detected proteins and the abundance of proteins enrich over 2-fold.

| **3D: Cells** | | | | | | | | | |
| --- | --- | --- | --- | --- | --- | --- | --- | --- | --- |
| **Cell culture** | **Total** | **2D** | **Only 2D** | **%** | **3D** | **Only 3D** | **%** | **3D vs 2D enrichment (2-fold)** | **%** |
| **Normoxic** | 7254 | 6669 | 278 | 3,832 % | 6976 | 585 | 8,065 % | 1020 | 14,061 % |
| **Hypoxic** | 7339 | 6769 | 271 | 3,693 % | 7068 | 570 | 7,767 % | 914 | 12,454 % |

Table S5. Comparison of protein amounts between 3D hypoxia and 2D normoxia in cells and EVs. Abbreviations: H = hypoxia, N = normoxia.

| **3D Hypoxic vs. 2D Normoxic** | | | | | | | | | |
| --- | --- | --- | --- | --- | --- | --- | --- | --- | --- |
| **Cell culture** | **Total** | **2D Normoxic** | **Only normoxic** | **%** | **3D hypoxic** | **Only hypoxic** | **%** | **3D H vs 2D N enrichment (2-fold)** | **%** |
| **Cells** | 7284 | 6669 | 216 | 2.965 % | 7068 | 615 | 8.443 % | 1023 | 14.044 % |
| **EVs** | 1199 | 666 | 74 | 6.172 % | 1125 | 559 | 46.622 % | 559 | 46.622 % |

Table S6. The impact of hypoxia on the number of detected proteins and the abundance of proteins enrich over 2-fold in EVs derived from 2D and 3D cell cultures. Abbreviations: H = hypoxia, N = normoxia.

| **Hypoxia: EVs** | | | | | | | | | |
| --- | --- | --- | --- | --- | --- | --- | --- | --- | --- |
| **Cell culture** | **Total** | **Normoxic** | **Only normoxic** | **%** | **Hypoxic** | **Only hypoxic** | **%** | **H vs N enrichment (2 fold)** | **%** |
| **2D** | 795 | 666 | 165 | 20.755 % | 630 | 129 | 16.226 % | 273 | 34.340 % |
| **3D** | 1344 | 1132 | 219 | 16.295 % | 1125 | 212 | 15.774 % | 280 | 20.833 % |

Table S7. The impact of 3D cell culture on the number of detected proteins and the enrichment of proteins in EVs derived from normoxic and hypoxic cell cultures.

| **3D: EVs** | | | | | | | | | |
| --- | --- | --- | --- | --- | --- | --- | --- | --- | --- |
| **Cell culture** | **Total** | **2D** | **Only 2D** | **%** | **3D** | **Only 3D** | **%** | **3D vs 2D enrichment (2-fold)** | **%** |
| **Normoxic** | 1194 | 666 | 62 | 5.193 % | 1132 | 528 | 44.221 % | 553 | 46.315 % |
| **Hypoxic** | 1207 | 630 | 82 | 6.794 % | 1125 | 577 | 47.804 % | 605 | 50.124 % |

Table S8.

| **Normoxia: EVs vs Cells** | | | | | | | | | |
| --- | --- | --- | --- | --- | --- | --- | --- | --- | --- |
|  | **Total** | **Cells** | **Cells only** | **%** | **EVs** | **EVs only** | **%** | **EV vs Cell enrichment (2-fold)** | **%** |
| **2D** | 6699 | 6669 | 6033 | 90,058 % | 666 | 30 | 0,448 % | 343 | 5,120 % |
| **3D** | 6984 | 6976 | 5852 | 83,792 % | 1132 | 8 | 0,115 % | 315 | 4,510 % |

Table S9.

| **Hypoxia: EVs vs Cells** | | | | | | | | | |
| --- | --- | --- | --- | --- | --- | --- | --- | --- | --- |
|  | **Total** | **Cells** | **Cells only** | **%** | **EVs** | **EVs only** | **%** | **EV vs Cell enrichment (2-fold)** | **%** |
| **2D** | 6833 | 6769 | 6203 | 99,063 % | 630 | 64 | 0,937 % | 371 | 5,430 % |
| **3D** | 7075 | 7068 | 5950 | 99,901 % | 1125 | 7 | 0,099 % | 298 | 4,212 % |

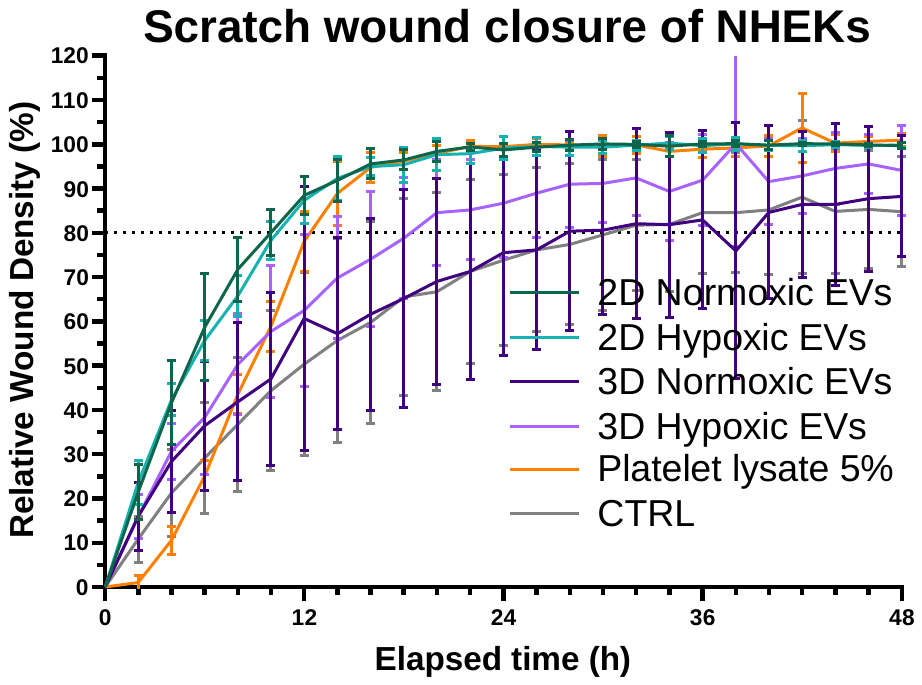

Figure S1. Therapeutic efficacy of MSC-derived EVs from hypoxic and 3D culture conditions in a 48 h NHEK scratch wound model. A: The impact of EVs on the closure time measured as relative wound density. Data presented as mean ± SD (N=4).

Table S10.

| Tukey's multiple comparisons test | Mean Diff, | 95,00% CI of diff, | Below threshold? | Summary | Adjusted P Value |  |
| --- | --- | --- | --- | --- | --- | --- |
| 2D Normoxic EVs vs. 2D Hypoxic EVs | 0,5573 | -0,3557 to 1,470 | No | ns | 0,4333 | A-B |
| 2D Normoxic EVs vs. 3D Normoxic EVs | 20,71 | 15,12 to 26,30 | Yes | **** | <0,0001 | A-C |
| 2D Normoxic EVs vs. 3D Hypoxic EVs | 11,73 | 7,247 to 16,20 | Yes | **** | <0,0001 | A-D |
| 2D Normoxic EVs vs. Platelet lysate 5% | 5,850 | -1,105 to 12,80 | No | ns | 0,1357 | A-F |
| 2D Normoxic EVs vs. CTRL | 22,46 | 16,39 to 28,53 | Yes | **** | <0,0001 | A-G |
| 2D Hypoxic EVs vs. 3D Normoxic EVs | 20,15 | 14,93 to 25,38 | Yes | **** | <0,0001 | B-C |
| 2D Hypoxic EVs vs. 3D Hypoxic EVs | 11,17 | 7,096 to 15,24 | Yes | **** | <0,0001 | B-D |
| 2D Hypoxic EVs vs. Platelet lysate 5% | 5,293 | -1,320 to 11,91 | No | ns | 0,1714 | B-F |
| 2D Hypoxic EVs vs. CTRL | 21,90 | 16,24 to 27,56 | Yes | **** | <0,0001 | B-G |
| 3D Normoxic EVs vs. 3D Hypoxic EVs | -8,986 | -12,35 to -5,624 | Yes | **** | <0,0001 | C-D |
| 3D Normoxic EVs vs. Platelet lysate 5% | -14,86 | -23,43 to -6,295 | Yes | *** | 0,0002 | C-F |
| 3D Normoxic EVs vs. CTRL | 1,750 | -0,4946 to 3,994 | No | ns | 0,1923 | C-G |
| 3D Hypoxic EVs vs. Platelet lysate 5% | -5,875 | -12,33 to 0,5838 | No | ns | 0,0896 | D-F |
| 3D Hypoxic EVs vs. CTRL | 10,74 | 8,272 to 13,20 | Yes | **** | <0,0001 | D-G |
| Platelet lysate 5% vs. CTRL | 16,61 | 8,899 to 24,32 | Yes | **** | <0,0001 | F-G |

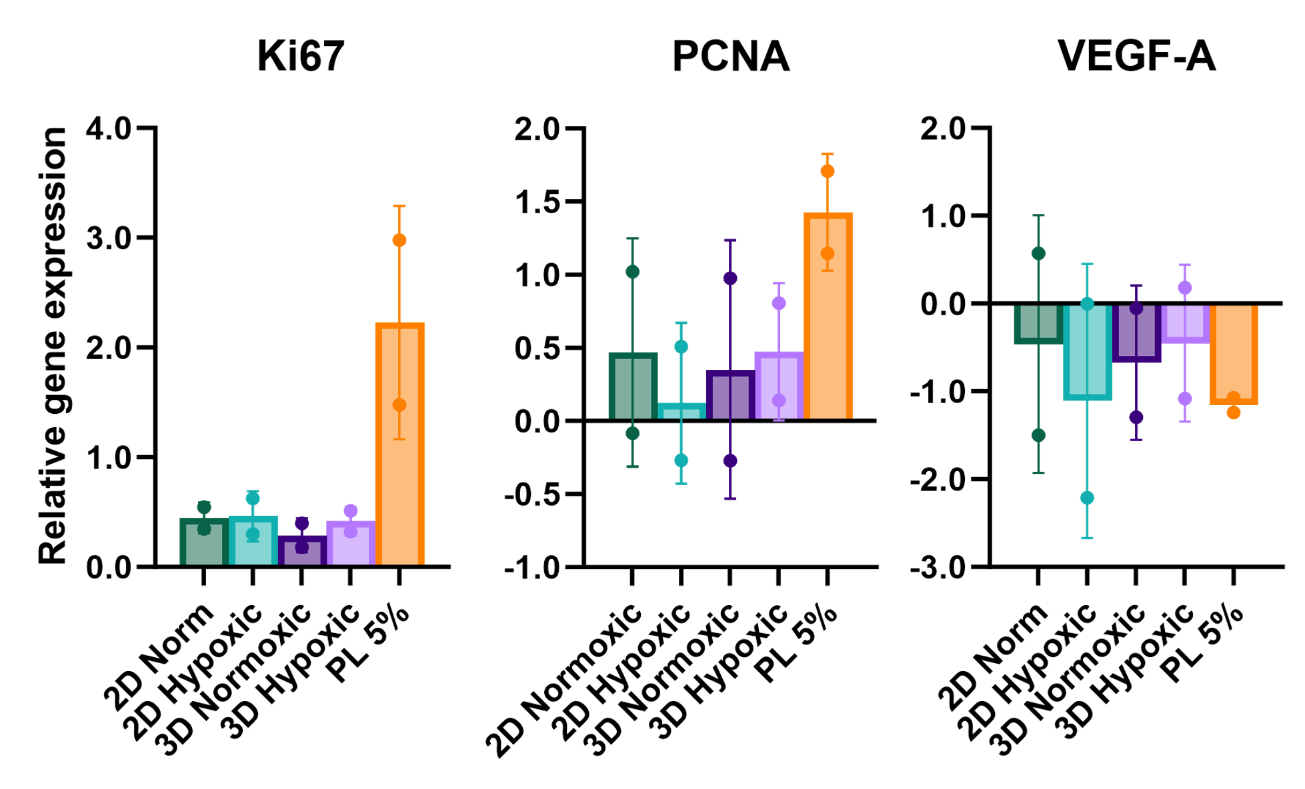

Figure S2. The fold change of relative gene expression of Ki67, PCNA and VEGF-A of NHEKs after 72 h scratch wound assay treated with MSC-derived EVs from hypoxic and 3D culture conditions. Data are represented as mean ± SD (N=2).
